# A synthetic cell-free pathway for biocatalytic upgrading of one-carbon substrates

**DOI:** 10.1101/2024.08.08.607227

**Authors:** Grant M. Landwehr, Bastian Vogeli, Cong Tian, Bharti Singal, Anika Gupta, Rebeca Lion, Edward H. Sargent, Ashty S. Karim, Michael C. Jewett

## Abstract

Biotechnological processes hold tremendous potential for the efficient and sustainable conversion of one-carbon (C1) substrates into complex multi-carbon products. However, the development of robust and versatile biocatalytic systems for this purpose remains a significant challenge. In this study, we report a hybrid electrochemical-biochemical cell-free system for the conversion of C1 substrates into the universal biological building block acetyl-CoA. The synthetic reductive formate pathway (ReForm) consists of five core enzymes catalyzing non-natural reactions that were established through a cell-free enzyme engineering platform. We demonstrate that ReForm works in a plug-and-play manner to accept diverse C1 substrates including CO_2_ equivalents. We anticipate that ReForm will facilitate efforts to build and improve synthetic C1 utilization pathways for a formate-based bioeconomy.

## Main

The accelerating climate crisis poses one of the most urgent economic and social challenges to humankind, driven by the unabated release and accumulation of CO_2_ in our atmosphere^1^. While significant strides have been made in commercializing carbon-free energy production, there remains a critical need for cost-effective, environmentally sustainable, and (cradle-to-gate) carbon-negative manufacturing of goods. An emerging solution lies at the intersection of chemistry and biology, where the electrochemical conversion of CO_2_ into soluble organic molecules provides substrates for enzymatic cascades to produce value-added chemicals^2,3^. Among these options, formate stands out as a promising bridge towards establishing a sustainable bioeconomy^4,5^. Formate can be efficiently generated through electrocatalysis^6^, exhibits high solubility in water, and simultaneously provides both a carbon source and reducing power. However, a significant hurdle exists—nature has only evolved a limited number of formate-fixing reactions, and, among those organisms discovered utilizing them, they are poorly suited for industrial applications^7^.

Recent efforts to develop a platform for formate utilization have focused on integrating formate assimilation pathways into workhorse biotechnology microbes such as *Escherichia coli*^8,9^ and *Cupriavidus necator*^10^. Despite notable progress, these synthetic formatotrophs have shown limited success in producing value-added chemicals, possibly due to unfavorable thermodynamic driving forces, environmental sensitivity, or inherent complexity of natural pathways^8,11^. Synthetic biology offers a potential way to address these challenges by designing new-to-nature metabolic pathways. When the design of such a system is freed from the evolutionary constraint to maintain life, it becomes possible to create pathways with superior thermodynamic or kinetic characteristics compared to those found in nature^3,12^.

Drawing inspiration from successful designer metabolic pathways^13–20^, we envisioned a synthetic formate assimilation pathway that is (i) thermodynamically favorable, (ii) tolerant to aerobic environments, (iii) composed of a minimal number of enzymes, (iv) independent from catalytic starting intermediates, and (v) devoid of a rate-limiting enzymatic CO_2_-fixing step by chemo-enzymatic coupling to electrochemically reduced CO_2_. A key requirement for our pathway architecture was the direct production of acetyl-CoA. Reaching this critical branchpoint between catabolism and anabolism unlocks access to an immense breadth of metabolic pathways that have been developed over the past decades to produce value-added chemicals^21^.

## Results

### Establishing a synthetic formate assimilation pathway

We designed a hypothetical, six-step reductive formate pathway (ReForm; **Fig. 1A**). ReForm was conceptualized around the engineered C1-C1 bond forming enzyme oxalyl-CoA decarboxylase (oxc), a variant of which was shown to catalyze the acyloin condensation between formyl-CoA and formaldehyde with a superior catalytic efficiency compared to similar C1-C1 bond forming enzymes (**Fig. S1-S3**)^22,23^. Our initial design starts with the activation of formate to formyl-CoA with an acyl-CoA synthetase (acs), followed by its reduction to formaldehyde with an acyl-CoA reductase (acr). Oxc ligates formyl-CoA and formaldehyde to form glycolyl-CoA, which is reduced to glycolaldehyde by an acr. Glycolaldehyde can then be dehydrated and phosphorylated to form acetyl-phosphate by a phosphoketolase (pk). A phosphotransacetylase (pta) then transfers a CoA onto acetyl-phosphate to form acetyl-CoA. A departure in our design philosophy from previous efforts was to build a completely synthetic pathway; that is, all five core reactions of the pathway are not known to readily occur in nature. Such synthetic designs expand the diversity of metabolic solutions for formate assimilation, offering a tailored fit to different host organisms or desired products.

**Figure 1.**
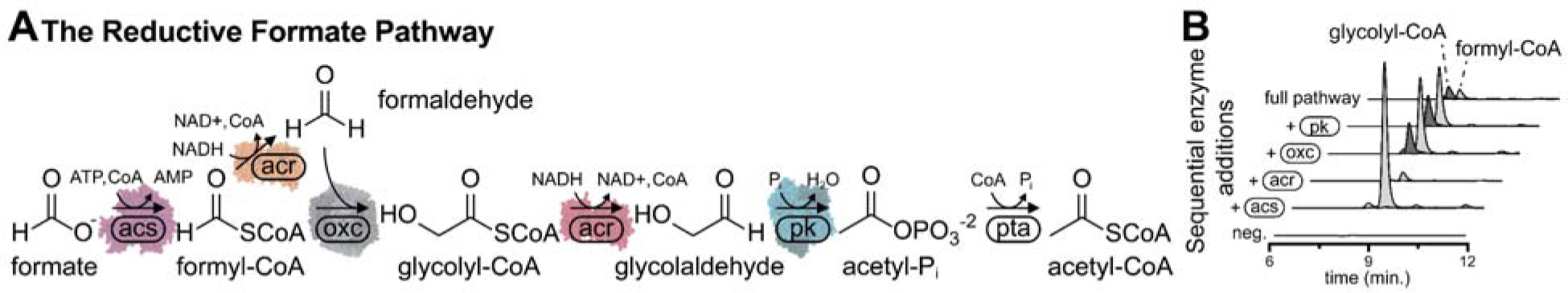
The synthetic reductive formate pathway. **(A)** Synthetic metabolic pathway to convert formate into acetyl-CoA, composed of five core new-to-nature reactions. **(B)** Each enzyme addition is labeled on the set of traces, starting with a no enzyme control. Extracted ion counts were determined for an *m/z* [M+H]^+^ of 797.1 for formyl-CoA, 828.1 for glycolyl-CoA, and 812.1 for acetyl-CoA, which corresponds to the mass of the CoA-thioester with the incorporation of 1 (formyl-CoA) or 2 (glycolyl-CoA and acetyl-CoA) ^13^C from ^13^C -formate. Traces are representative of *n* = 3 independent reactions.

ReForm consists of six individual enzyme-catalyzed reactions, with five core reactions being new-to-nature. We began by searching the literature for enzymes with a demonstrated capacity to catalyze these reactions based on promiscuity (i.e., a given enzyme can perform similar chemistries on molecules similar to the native substrate). We selected an acs from *Erythrobacter* sp. NAP1 which natively transforms acetate to acetyl-CoA (instead of formate to formyl-CoA)^24^, an acr from *R. palustris* which natively reduces propionyl-CoA to propionaldehyde (instead of reducing formyl-CoA and glycolyl-CoA)^25^, and a pk from *B. adolescentis*, which natively cleaves D-fructose 6-Pi into acetyl-Pi and erythrose 4-Pi (instead of glycolaldehyde into acetyl-Pi)^16^.

To evaluate the feasibility of ReForm, we carried out a stepwise construction of the pathway by sequentially adding each purified enzyme to a buffer containing a labeled ^13^C-isotope of formate and necessary cofactors. HPLC-MS analysis of all three CoA-thioesters demonstrated that the initial sequence of the pathway was functional, but we did not observe the production of acetyl-CoA from the complete pathway (**Fig. 1B** and **Fig. S4**). We hypothesized that the promiscuous acyl-CoA reductase (acr) from *R. palustris* could reduce detection of acetyl-CoA through the unwanted reduction of acetyl-CoA to acetaldehyde^26^ (**Fig. S5**). To sequester any synthesized acetyl-CoA, an additional malate read-out module was created by the addition of a malate synthase (mls) from *E. coli*^15^. While we were able to detect small quantities of malate (∼1 µM; **Fig. S6**), the selected enzymes’ low activities with non-native substrates presented a major bottleneck.

### Engineering a phosphoketolase into an acetyl-P_i_ synthase

To improve the titer and rate of our pathway, we sought to engineer the enzymes of ReForm. Our general strategy was to search for homologous enzymes that have high promiscuity towards the desired reactions and then apply directed evolution principles to engineer the enzymes for greater activities with the non-native substrates. A key feature of our approach was the use of cell-free gene expression (CFE) systems for the rapid synthesis and functional testing of both protein homologs and mutants^27–29^. We demonstrated this approach by first targeting the phosphoketolase reaction for producing acetyl-P_i_, given its demonstrated low native activity for this reaction (∼0.005 µM min^-1^ mg^-1^)^16,30^ (**Fig. 2a**). We constructed a curated phylogenetic tree of the entire annotated pk protein family (IPR005593, comprised of 15k members; **Fig. S7**), of which we selected a diverse set of 30 homologs to express, purify, and test for activity with glycolaldehyde (**Fig. 2b** and **Fig. S8**).

**Figure 2.**
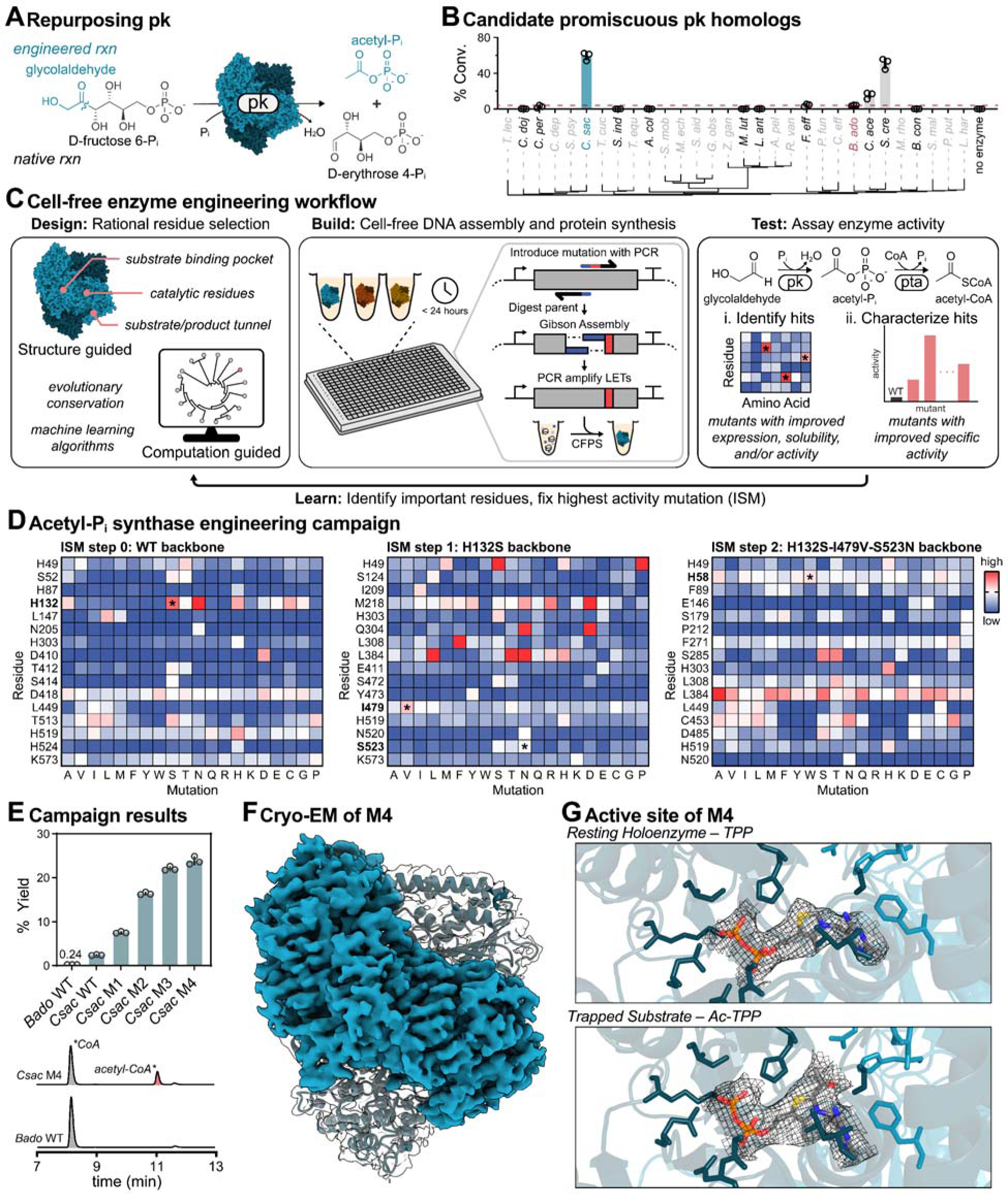
Rational engineering of a phosphoketolase into an acetyl-P_i_ synthase. **(A)** The native phosphorolytic cleavage activity of pk to produce acetyl-P_i_ resembles the target engineered reaction. **(B)** Exploring a diverse set of homologs resulted in several enzymes with activity for glycolaldehyde. Percent conversion was determined by quantification of CoA and acetyl-CoA (*n* = 3, error bars indicate ± SD). **(C)** A schematic of the cell-free protein engineering workflow. Site saturation mutagenesis and cell-free protein expression are carried out in less than 24 hours to generate sequence-defined libraries. **(D)** Iterative site saturation mutagenesis of 16 residues showing percent conversion normalized to the step backbone (*n* = 1). After further characterization, four substitutions (H58W, H132S, I479V, and S523N; asterisks) were identified and resulted in a quadruple mutant. **(E)** HPLC traces of acetyl-CoA product (red) and CoA substrate (purple) of wild-type pk and engineered mutants. Traces are single samples representative of *n* = 3 independent experiments. **(F)** Electron density map and model of *Csac* M4 complexed with Ac-TPP at a global resolution of 2.30 Å (PDB: 9CD4). **(G)** Model of the active site of *Csac* M4 complexed with TPP (PDB: 9CD3) and Ac-TPP (PDB: 9CD4) overlayed with their respective electron density map of the cofactor.

We identified several homologs with a higher activity than our initial candidate enzyme from literature, notably one (*Csac* Pk) with an approximately 10-fold increase under the tested *in vitro* reaction conditions. Based on structural considerations (i.e., 6 Å around the carbanion of the thiamine diphosphate cofactor ylid), evolutionary conservation (i.e., an EVmutation probability density model trained on a multiple sequence alignment of evolutionarily related sequences)^31^, and a deep learning model trained to optimize local amino acid microenvironments^32^, we initially selected 16 residues potentially important for catalysis and substrate specificity to mutate (**Fig. 2c**). We used iterative site saturation mutagenesis (ISM) to identify and accumulate beneficial mutations of these residues for acetyl-P_i_ synthesis (**Fig. S9** and **Fig. S10**). After three rounds of ISM and exploring 1,200 unique, sequence-defined mutants among 37 residues, we obtained a quadruple mutant (*Csac* M4) with a more than 10-fold increase in catalytic efficiency compared to wild-type (k_cat_/K_M_ increase from 0.085 ± 0.016 to 1.18 ± 0.17 mM^-1^ min^-1^, respectively) (**Fig. 2d-e** and **Fig. S11**).

To gain insights into the topology of the engineered enzyme’s active site, we solved the structure of *Csac* M4 at a resolution of 2.30 Å with Cryo-EM (**Fig. 2F** and **Fig. S12-S13**). Soaking the enzyme with glycolaldehyde without the addition of phosphate as the resolving nucleophile enabled us to gain a glimpse of the reaction mechanism by covalently trapping glycolaldehyde on TPP (**Fig. 2G**). This reaction intermediate, 2-acetyl-TPP (Ac-TPP) is shared between the canonical reaction mechanism and the engineered enzyme, enabling the switch in catalysis from phosphorlytic cleavage of sugars to the phosphorylation and dehydration of glycolaldehyde^16,33^. Observing the Ac-TPP cofactor provides insight into the non-native reactivity with glycolaldehyde and could help guide future pk engineering efforts for sustainability applications^30^.

### Divergent directed evolution of substrate-specific acyl-CoA reductases

After having success with our phosphoketolase engineering campaign, we next targeted the two acyl-CoA reductase reactions of ReForm. Engineering acr presented a unique challenge in that we needed to decrease activity with acetyl-CoA while simultaneously improving activity for formyl-CoA and glycolyl-CoA. To help control specificity, our approach was to engineer two distinct enzymes to catalyze these respective reductions instead of a single bifunctional enzyme. We selected 34 candidates from among three protein families with reported acylating dehydrogenase activity (IPR013357, IPR012408, IPR00361; **Fig. S14**) to express and characterize (**Fig. S15**). Serendipitously, we identified a single acr with activity (∼2-fold increase above *Rpal* Acr) for both formyl-CoA (acr_f_) and glycolyl-CoA (acr_g_) from the anaerobic photoautotrophic bacteria *C. thalassium* (**Fig. S16**). In our first round of ISM, we screened 16 residues against all three CoA-thioesters (915 unique substrate-mutant pair reactions) to find mutations that selectively increased activity for formyl-CoA or glycolyl-CoA and/or decreased activity for acetyl-CoA (**Fig. S17**). The evolution paths bifurcated in subsequent rounds of mutagenesis after we identified different beneficial mutations in the first round for each substrate (**Fig. S18** and **S19**). Subsequent rounds of ISM focused on a down-selected set of 8 residues identified in the first round to impact substrate preference. We identified an acr_f_ triple mutant (*Cthal_f_* M3, **Fig. 3A**) with a 15-fold increase in specificity and an acr_g_ double mutant (*Cthal*_g_ M2, **Fig. 3B**) with a 13-fold increase in specificity (**Fig. S20**). While all mutations are found in the substrate binding pocket, the final mutants notably have no mutated residues in common (**Fig. S21**).

**Figure 3.**
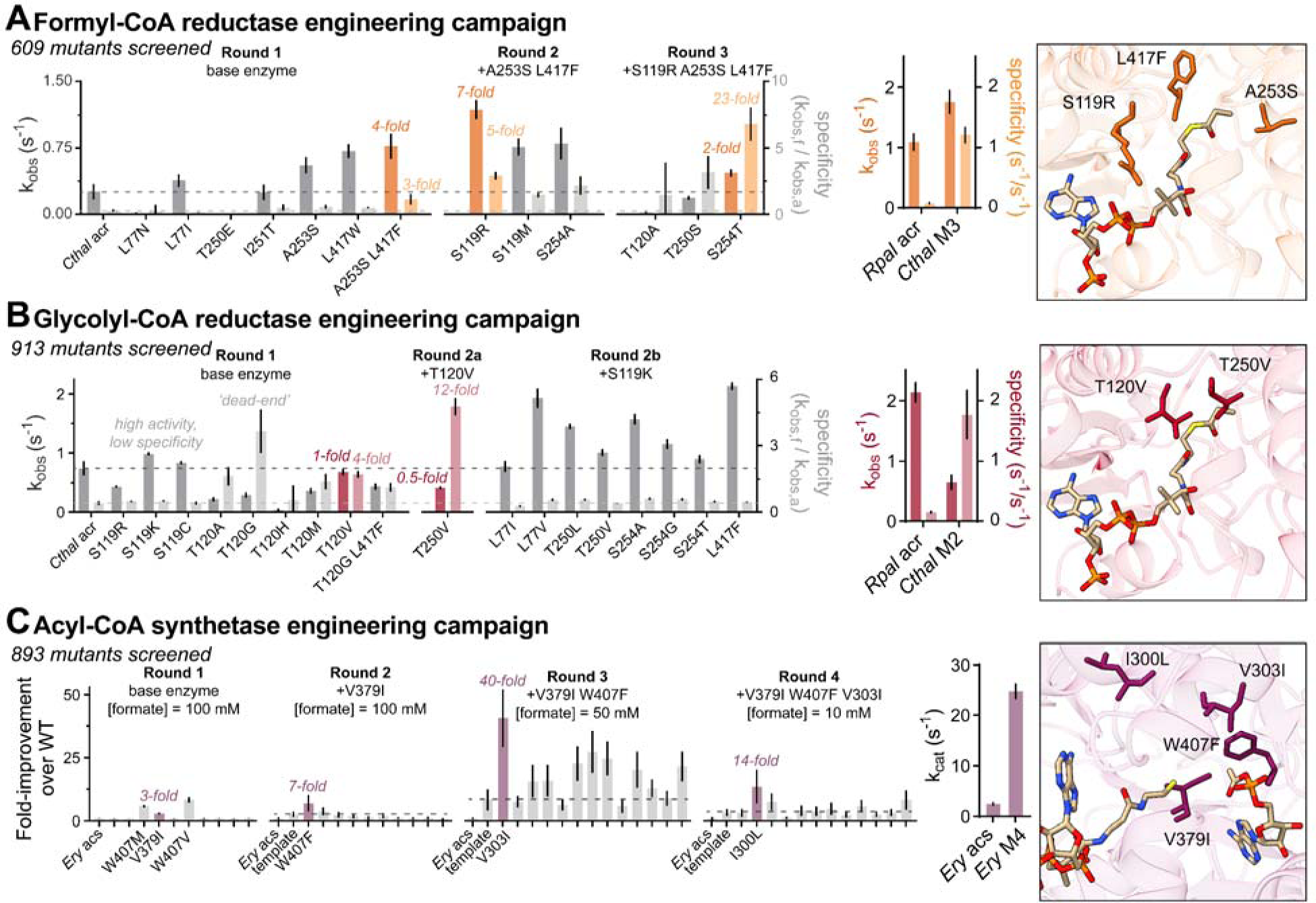
Enzyme engineering campaigns and characterized mutants for the (A) formyl-CoA reductase, (B) glycolyl-CoA reductase, and (C) acyl-CoA synthetase reactions. The mutant showing the best parameters that was carried forward in each round is highlighted. Campaigns resulted in remodeled active sites to accept non-native substrates, with the residues that were mutated labeled. All values are either from *n = 3* independent measurements or are results from Michaelis-Menten fitting from *n* = 3 independent experiments. Complete engineering campaigns can be found in the SI.

### Engineering a formyl-CoA synthetase

Our final enzyme engineering challenge was the initial acyl-CoA synthetase reaction to activate formate. Given that formate would be exogenously supplied to the reaction in saturating quantities, we focused on improving the turnover number (k_cat_) of acs. Our starting point was an acetate-activating acs from *Erythrobacter* sp. NAP1, which was successfully engineered to change its native substrate specificity to glycolate, highlighting its engineerability^24^. Exploring 18 residues over four rounds of ISM resulted in a quadruple mutant (*Enap* M4) with a 2-fold increase in k_cat_ for formate (**Fig. S22-S24**). All four accumulated mutations were in the putative substrate binding pocket and fill in the active site with bulky, hydrophobic amino acids (**Fig. S25**). Expression of acs in a strain of *E. coli* with a genomic knockout of a lysine acetyltransferase that post-translationally inactivates acs *in vivo* further increased k_cat_ by another 5-fold, resulting in a 10-fold overall increase in activity (2.5 ± 0.3 to 25 ± 1.7 s^-1^; **Fig. 3C** and **Fig. S24**)^24,34^. While we did not observe a significant shift in *K*_M_ throughout the engineering campaign, the measured values do fall in similar ranges to naturally found formate activating enzymes (e.g., formate-tetrahydrofolate ligase^35^), indicating we may have reached the inherent natural limit to formate affinity given its low molecular mass and low hydrophobicity^36^.

### ReForm excels at assimilating diverse C1 substrates

Taken together, our comprehensive ISM campaigns assessed 3,173 sequence-defined enzyme variants to identify four enzyme variants for the five, core new-to-nature reactions in ReForm (**Fig. 3b**, **Fig. 3c**, and **Fig. S26)**. To increase titers (**Fig. S27**), we carried out an optimization of the pathway in three steps. First, we tuned cofactor and enzyme loading with definitive screening design (DSD; **Fig. S28**). The objective was strictly to increase malate titers given a range of possible cofactor and enzyme concentrations. We restricted maximum values of all components to prevent superior conditions resulting simply from increase enzyme loading (enzyme concentrations were capped between 1-20 µM). Second, we added cofactor regeneration to recycle cofactors and keep concentrations high (**Fig. S29**). A formate dehydrogenase was added to regenerate NADH from formate while a polyphosphate kinase and polyphosphate were added to regenerate ATP^37^. Third, we added a formyl-phosphate reductase to recycle formyl-phosphate that could be produced from an unwanted side-reaction between formyl-CoA and the phosphotransacetylase (**Fig. S30**)^20^. When combined, ReForm pathway titers increased by three orders of magnitude over our original pathway iteration (maximum titer of 318 ± 3 µM of malate at 24 hours; **Fig. 4a-b** and **Fig. S31**).

**Figure 4.**
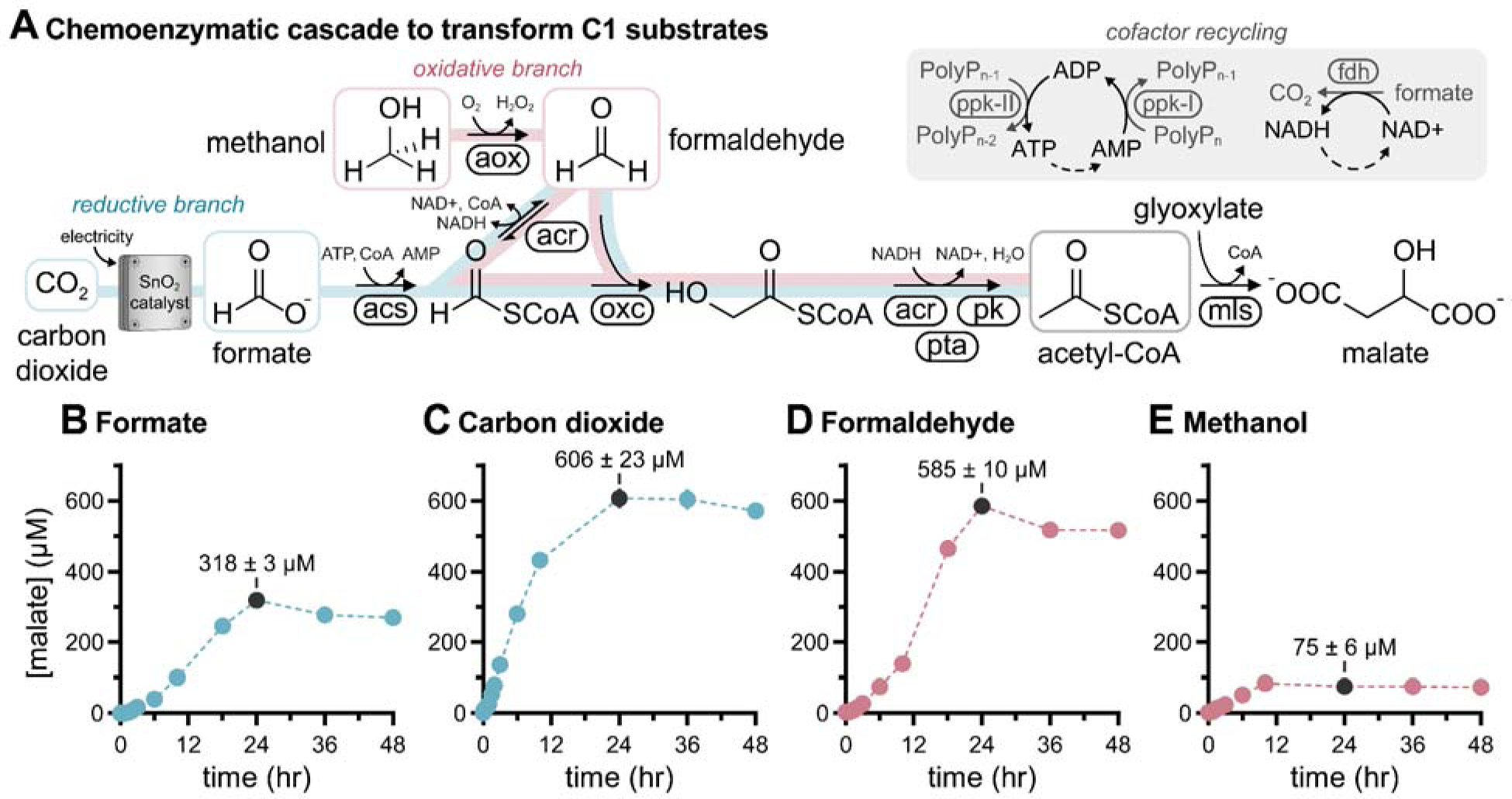
Chemoenzymatic conversion of C1 substrates into acetyl-CoA. **(A)** The combined chemoenzymatic cascade enables the assimilation of C1 substrates (formate, CO_2_, formaldehyde, and methanol) into acetyl-CoA. Malate synthase (mls) is added to convert acetyl-CoA into malate. **(B)** Time course of malate synthesis with formate as a substrate. **(C)** Time course of malate synthesis with formate derived from the electrochemical reduction of CO_2_ as a substrate. **(D)** The maximum observed malate titer for each substrate measured at 24 hours for formate, CO_2_, and formaldehyde and 10 hours for methanol. **(B-D)** Malate concentration was determined by LC-MS with an *m/z* [M-H]^-^ of 133.01 for ^12^C malate (CO_2_, formaldehyde, and methanol as substrates) and 135.01 for the malate containing two ^13^C from ^13^C-formate (formate as a substrate) (*n* = 3, error bars indicate ± SD).

We next attempted to synthesize acetyl-CoA directly from CO_2_ by coupling ReForm with the electrochemical reduction of CO_2_ using a commercial SnO_2_ inorganic catalyst. Formate was initially produced in isolation in the electrochemical module at a rate of 150 mM hr^-1^ mg^-1^ to a concentration of 50 mM within 20 minutes of electrosynthesis and confirmed by NMR (**Fig. S32**). Following the direct addition of the crude formate to the enzymatic formate assimilation module, we observed even higher maximum titers of malate than the standalone enzymatic module (606 ± 23 µM of malate at 24 hours; **Fig. 4c** and **Fig. S33**).

With the demonstrated success of ReForm to produce acetyl-CoA from both formate and CO_2_, we wondered if it was possible to additionally assimilate other relevant C1 substrates. We designed routes through ReForm to convert both formaldehyde and methanol into acetyl-CoA with no changes to the core architecture of the pathway except for simply removing the first enzymatic step (acs; **Fig. 4a**). Notably, these two variations are completely cofactor neutral (i.e., there is no net theoretical change in reducing equivalents or ATP per molecule of acetyl-CoA produced). Following pathway optimization, we were able to reach similar titers of acetyl-CoA with formate as a substrate (maximum titers of 585 ± 10 µM and 84 ± 2 µM of malate at 24 hours from formaldehyde and methanol, respectively; **Fig. 4d** and **Fig. S34-S36**). The tradeoff of utilizing methanol and formaldehyde as substrates is that, while possessing a higher thermodynamic driving force, an additional source of electrons is needed to provide reducing power for reactions beyond acetyl-CoA that require them.

## Discussion

In this work, we demonstrate a synthetic chemoenzymatic cascade called ReForm towards the valorization of organic and inorganic C1-substrates. The five new-to-nature reactions of ReForm were selected among 66 enzyme candidates from both eukaryotic and prokaryotic origins, extensively engineered by testing 3,173 sequence-defined mutants, and combined in a cell-free, plug-and-play manner to convert diverse C1-substrates into acetyl-CoA. Through enzyme engineering and pathway optimization, we were able to improve malate titers by several orders of magnitude compared to the initial pathway design when utilizing CO_2_ as a carbon source, from limit of detection to ∼600 µM. The successful demonstration of this system paves the way for future applications, including the *in vitro* synthesis of value-added chemicals such as biofuels and terpenes directly from CO_2_^38,39^.

Notably, ReForm consisting of only six reactions affords a lower total enzyme loading to achieve higher possible specific titers as compared to longer synthetic pathways (∼4 mg/mL total loading; less than half of the state-of-the-art *in vitro* synthetic pathway THETA)^14^. The energy efficiency of ReForm (only 4 ATP and 2 NADH equivalents per acetyl-CoA compared to 7 ATP and 4 NAD(P)H of the Calvin Cycle and 4 ATP and 5 NAD(P)H of the THETA cycle; **Fig. S37**) helps enable similar titers of acetyl-CoA to be achieved while operating at these lower loadings (**Table S1**)^14,40^. While excelling in specific titers and reaction longevity as compared to the THETA and CETCH cycles^40^, ReForm has lower peak productivities (**Table S1**) – a potential trade-off caused by removing catalytic starter molecules seen in cyclic pathways. However, the electrochemical production of formate additionally provides the dual advantage of carbon coming from CO_2_ and electrons coming from electricity, eliminating the need for a sacrificial substrate to provide reducing power as seen in the CETCH and THETA cycles. The linear nature and minimal overlap with central metabolism of ReForm may also simplify its implementation into living cells^14^. However, to achieve economic relevance, improvements in volumetric productivities (g product L^-1^ hr^-1^), reaction longevity, and total turnover per catalyst must still be made.

By combining electrochemistry and synthetic biology, our work expands the possible solution space of generalizable CO_2_-fixation strategies. While these strategies have been successful in their own right^41,42^, we anticipate their partnership will become critical to provide a carbon-negative alternative to traditional chemical synthesis in the future.

## Methods

### Materials and bacterial strains

All consumables were purchased from Sigma-Aldrich unless stated otherwise. Standard microtiter plates (96- and 384-well) were purchased from BioRad. ^13^C-sodium formate was purchased from Cambridge Isotope Laboratories Inc. (MA, USA). DNA for all enzymes used was ordered from Twist Bioscience (CA, USA) in the vectors pJL1 (Addgene #69496) for expression via CFPS or pETBCS, a modified pET-22b vector^43^ (Novagen/EMD Millipore, Darmstadt, Germany) for recombinant expression in *E. coli*. Codon optimization for *E. coli* was either performed using Integrated DNA Technologies (IDT) or Twist Bioscience. NEB® 5-alpha chemically competent *E. coli* cells were used for cloning (NEB), BL21 Star™ (DE3) (Invitrogen) cells were used for cell-free lysate production, and BL21 (DE3) chemically competent cells (NEB) were used for recombinant expression of proteins. For the expression of acyl-CoA synthetases, an *E. coli* BL21 (DE3) *patZ* knockout was created using a CRISPR-Cas9 and λ-red recombination-based method^44,45^. Briefly, the CRISPR endonuclease introduces a double-stranded DNA break in a locus of interest. Provided with a donor DNA that is homologous to sections of the chromosome on 5’ and 3’ ends but lacks the knockout gene, λ-red proteins will recombine the donor DNA into the chromosome. This simultaneously removes the gene of interest and the site where the DNA break is occurring, allowing the cells to survive if the knockout was successful. To confirm the knockout of *patZ*, single colonies were picked and analyzed by colony PCR with primers flanking the gene.

### Cell-free library generation and protein synthesis

Crude cell extracts were prepared as previously described using *E. coli* BL21 Star (DE3) cells (Invitrogen)^46^. CFPS reactions were performed based on the Cytomim system^47,48^ and, unless otherwise noted, carried out in 96-well or 384-well PCR plates (Bio-Rad) as 15-µL reactions with 1 µL of linear expression template (LET) serving as the DNA template. Briefly, 15-μL reactions were carried out with final concentrations of 8 mM magnesium glutamate, 10 mM ammonium glutamate, 130 mM potassium glutamate, 1.2 mM ATP, 0.85 mM of GTP, CTP, and UTP each, 0.03 mg/mL folinic acid, 0.17 mg/mL tRNA (Roche), 0.4 mM NAD+, 0.27 mM CoA, 4 mM oxalic acid, 1 mM putrescine, 1.5 mM spermidine, 57 mM HEPES pH 7.2, 33 mM phosphoenolpyruvate (Roche), 30% v/v cell extract, and remaining volume with water to 15 μL. Reactions were incubated at 30 °C for 16-20 hours.

Cell-free site saturation mutagenesis was performed as described previously. Briefly, primers were designed using Benchling with melting temperature calculated by the default SantaLucia 1998 algorithm. The general heuristics we followed for primer design were a reverse primer of 58 °C, a forward primer of 62 °C, and a homologous overlap of approximately 45 °C. All primers were ordered from IDT; forward primers were synthesized in 384-well plates normalized to 2-µM for ease of setting up reactions. All cloning steps were set up using an Integra VIAFLO liquid handling robot in 384-well PCR plates (Bio-Rad). The cell-free library generation was performed as follows: (1) the first PCR was performed in a 10 µL reaction with 1 ng of plasmid template added, (2) 1 µL of DpnI (NEB) was added and incubated at 37°C for two hours, (3) the PCR was diluted 1:4 by the addition of 29 µL of nuclease-free (NF) water, (4) 1 µL of diluted DNA was added to a 3 µL Gibson assembly reaction and incubated for 50 °C for one hour, (5) the assembly reaction was diluted 1:10 by the addition of 36 µL of NF water, (6) 1 µL of the diluted assembly reaction was added to a 9 µL PCR reaction. All PCR reactions used Q5 Hot Start DNA Polymerase (NEB). The product of the second PCR is a LET for expression in CFPS, which are amplified using universal forward (CTGAGATACCTACAGCGTGAGC) and reverse (CGTCACTCATGGTGATTTCTCACTTG) primers. To accumulate mutations for ISM, 3 µL of the round “winner” from the diluted Gibson assembly plate was transformed into 20 µL of chemically competent *E. coli* (NEB 5-alpha cells). Cells were plated onto LB plates containing 50 µg/mL kanamycin (LB-Kan). A single colony was used to inoculate a 50 mL overnight culture of LB-Kan, grown at 37 °C with 250 RPM shaking. The plasmid was purified using ZymoPURE II Midiprep kits (Zymo Research) and sequence confirmed.

### Cryo-EM sample preparation and data collection

*Csac* Pk M4 was purified as described below. The protein was then additionally purified to homogeneity using a Cytiva HiPrep™ Sephacryl™ S-200 HR Column according to the manufacturer’s specifications with a running buffer containing 50 mM HEPES pH 7.4, 150 mM NaCl, and 250 µM MgCl_2_. After purification, thiamine pyrophosphate (TPP) was added to a final concentration of 50 µM. To covalently trap glycolaldehyde on TPP, 1 mg/mL of purified protein was soaked in 25 mM glycolaldehyde for 10 minutes at room temperature and then the buffer was exchanged into the running buffer using an Amicon Ultra-0.5 centrifugal filters (50 kDa MWCO; EMD Millipore).

Ultrafoil 1.2/1.3 Au 300 mesh grids were glow discharged and 3ul of the protein sample at 1 mg/ml was applied to the grid surface. The grids were incubated for 5 sec in the humidity chamber of Vitrobot Mark IV set at 100% humidity and 4 degrees Celsius. Grids were blotted at blot force 2 for 3 seconds and quickly plunged frozen in liquid ethane. Cryo-EM imaging was performed on a ThermoFisher Titan Krios equipped with Falcon 4i direct electron detector and SelectrisX post-column energy filter. The microscope was operated at 300 kV accelerating voltage with a nominal magnification of x130000 that resulted in a magnified pixel size of 0.98 A. Each movie was recorded at the total electron dose of 40e-/A2 over 38 frames. The images were obtained at the defocus ranging from -0.8 to -1.8um.

### Image processing and 3D reconstruction

Preprocessing of both datasets was performed using cryoSPARC^49^. Dose fractionated movies were subjected to beam-induced motion correction and dose weighting using Patch motion correction. Contrast transfer function (CTF) parameters were performed for each motion-corrected micrograph followed by micrographs curation based on parameters such as average intensity and CTF fit resolution. 3,159,785 particles were extracted from 5900 micrographs for the *Csac* Pk M4-TPP and 3,761,029 particles were extracted from 6652 micrographs for the *Csac* Pk M4-HDE/HTL dataset. Subsequently, 3-5 rounds of 2d classification were performed followed by multi class ab initio and heterogenous refinement. Homogenous refinement was performed on the largest class from the heterogeneously refinement job on the binned particles. Further, the particles were re-extracted to 0.95 Å/pix, and another round of homogenous refinement was performed. UCSF ChimeraX (v.1.7)^50^ was used for map and model visualization.

### Molecular model building

AlphaFold2 was used to generate a model dimer of *Csac* Pk M4 using its protein sequence^51^. The X ray crystal structure of phosphoketolase from *Bifidobacterium breve* complexed with TPP/HDE/HTL (PDB: 3ahc, 3ahe, and 3ahd)^52^ was used to align with the AlphaFold model to roughly estimate the position of ligand in the maps. The ligands were combined with the AphaFold-generated dimer model and were fitted into the maps using the ChimeraX ‘fit-in-map’ function. To improve the modelling, several rounds of interactive model adjustment in Coot (v0.9.8.8 EL) followed by Real-space refinement of the fitted model was performed in Phenix (v 1.21.1-5286) from SbGrid suite^53^ using secondary structure restraints in addition to default restraints. The final model was generated using Phenix refinement.

### ReForm Assays

All reactions were performed at 30 °C with reaction size noted below. All reactions were quenched by addition of 5 µL 10% w/v formic acid to 20 µL of sample (or 3.75 µL 10% w/v formic acid to 15 µL of sample), centrifuged at 4,000 x *g* for 10 min at 4 °C to remove precipitated protein, and either immediately prepared for analysis or stored at -80 °C until needed. To prepare for MS analysis of CoA-thioesters and malate, 20 µL of quenched reaction was transferred to a clean vial and diluted 1:2 with 20 µL of H_2_O.

#### Stepwise pathway construction

Final reactions contained 20 mM sodium phosphate buffer pH 7.4, 5 mM ATP, 6.5 mM NADH, 10 mM MgCl_2_, 0.5 mM CoA, 50 µM TPP, 1 mM glyoxylate, 50 mM ^13^C sodium formate, 5 U/mL pyrophosphatase (Sigma-Aldrich I5907), 3 µM *Enap* Acs, 3 µM *Rpal* PduP, 10 µM *Mext* Oxc M4, 10 µM *Bado* Pk, 0.25 µM *Ecol* Pta, and 1 µM *Ecol* Mls. A reagent mix was first made with all components except for the core ReForm enzymes and distributed into seven 1.5 mL Eppendorf tubes. Enzymes were added to the final concentrations listed above with each sequential reaction containing an additional enzyme. Extra volume of H_2_O was added to bring up the final reaction volume to 200 µL. A negative control containing only H_2_O and no additional enzyme was also included. These reactions were sampled for both CoA-thioesters and malate. 20-µL samples were taken at various time intervals (10, 30, 60, 90, 120, 150, 180, and 240 minutes) to ensure we would not miss any time resolved production of intermediates or malate.

Initial unsuccessful pathway attempts (that produced detectable formyl-CoA and glycolyl-CoA but no detectable acetyl-CoA or malate) utilized lower concentrations of *Bado* Pk (5 µM) and *Mext* Oxc M4 (5 µM), lower concentrations of *Enap* Acs and *Rpal* PduP (1 µM each), and higher concentrations of *Ecol* Pta (1 µM). Pta was decreased after observing potential side reactivity with formyl-CoA (an observed decrease in formyl-CoA after the direct addition of pta).

#### Leave one out

Reactions were set up as described for the stepwise pathway construction except for the decrease of *Ecol* Pta from 0.25 µM to 0.125 µM. A reagent mix was first made with all components except for the core ReForm enzymes and formate and distributed into eight 1.5 mL Eppendorf tubes. Each reaction had a single enzyme removed (or formate) with extra volume of H_2_O added to bring up the final reaction volume to 90 µL. 20-µL samples were taken at various time intervals (60, 120, 180, and 240 minutes) to ensure we would not miss any time resolved production of intermediates or malate.

#### Definitive screening design (DSD)

The 3-level screening design was performed using JMP Pro 17, with upper and lower limits and the reaction components varied found in the corresponding SI Figure. All engineered variants of enzymes were used in this experiment. We first performed a screen of 21 reaction conditions to map out the relationship between all tested variables and malate production. Final reactions contained 20 mM sodium phosphate buffer pH 7.4, 2 mM ATP, 4 mM NADH, 10 mM MgCl_2_, 50 µM TPP, 1 mM glyoxylate, 50 mM ^13^C sodium formate, 5 U/mL pyrophosphatase (Sigma-Aldrich I5907), and 0.5 µM *Ecol* Mls. The concentrations of remaining enzymes and CoA varied between each reaction condition. A reagent mix was first made with all components except for the varied enzymes and CoA and distributed into a 96-well PCR plate. Different dilutions of each varied component were added to the reactions to bring up the total volume to 15 µL, with each condition ran in triplicate for four hours. With the resulting data, three regression models were fit with default variables: stepwise, stepwise (exclusive to non-zero data points), and support vector machines (SVM). All models were fit using default parameters that only modeled first-order interactions between variables. For stepwise models, a minimum BIC stopping rule was employed. To determine predicted optimized conditions, the prediction profiler was set to maximize desirability. We then went on to validate predictions by comparing the best condition that was observed in the screen with the three predicted conditions. The stepwise prediction set everything to the upper bound of the DSD screen: 1 mM CoA, 2 µM acs, 5 µM acr_f_, 5 µM acr_g_, 20 µM oxc, 20 µM pk, and 0.01 µM pta. The stepwise (non-zero) prediction was identical except set oxc to the lower bound of the DSD screen (2 µM). The SVM prediction had 0.94 mM CoA, 1.18 µM acs, 3.63 µM acr_f_, 3.66 µM acr_g_, 14.27 µM oxc, 20 µM pk, and 0.43 µM pta. These reactions were set up identically to the DSD screen.

After optimizing the pathway for malate production from formate, we repeated DSD with the variant of ReForm that uses formaldehyde as a substrate. As above, all engineered enzymes were used in this experiment and the reaction components varied can be found in the corresponding SI Figure. We first performed a screen of 21 reaction conditions to map out the relationship between all tested variables and malate production. Final reactions contained 20 mM sodium phosphate buffer pH 7.4, 10 mM MgCl_2_, 50 µM TPP, 1 mM glyoxylate, 50 mM formaldehyde, and 0.5 µM *Ecol* Mls. The concentrations of remaining enzymes, CoA, and NAD+ varied between each reaction condition. Three models were then fit like above, however, as there was no measured zero malate condition, we only included a single stepwise model. The stepwise optimized conditions were 0.2 mM CoA, 0.5 mM NAD+, 5 µM acr_f_, 5 µM acr_g_, 2 µM oxc, 20 µM pk, and 0.01 µM pta. The SVM optimized conditions were 0.78 mM CoA, 2.19 mM NAD+, 5 µM acr_f_, 5 µM acr_g_, 15.5 µM oxc, 20 µM pk, and 0.384 µM pta. These reactions were set up identically to the DSD screen, and an additional experiment was ran using 20 mM formaldehyde to examine potential inhibitory effects.

#### Cofactor regeneration

Using the DSD optimized conditions with formate as a substrate, we then added cofactor regeneration enzymes for ATP and NADH. Final reactions conditions were the same as above with two additional polyphosphate kinases (*Rmel* ppk1 and *Ajoh* ppk2) added at 0.4 or 2 µM and an additional formate dehydrogenase (*P101* fdh) added at 1 or 6 µM. Polyphosphate (adjusted to pH 7 with KOH) was added at a final concentration of 10 mM (phosphate equivalents) and ^13^C sodium formate was increased to 100 mM. Reactions were performed in triplicate at 15 µL and incubated for four hours.

#### Metabolic proofreading

After determining cofactor regeneration enzyme conditions, we wanted to test if adding in a recently engineered formyl-phosphate reductase (fpr), DaArgC3, could improve pathway activity^20^. Earlier experiments suggested that pta could be exhibiting potential side activity with formyl-CoA and producing formyl-P_i_, a dead-end metabolite for ReForm. We hypothesized that adding in fpr could reduce the produced formyl-P_i_ into formaldehyde to introduce the wasted metabolite back into the pathway. Using the conditions from the cofactor regeneration screen (2 µM of ppk1 and ppk2 and 1 µM of fdh) as the base reaction, we added 0.5, 1 or 5 µM of fpr. Reactions were performed in triplicate at 15 µL and incubated for four hours.

#### Time course

The metabolic proofreading strategy did not provide any quantifiable benefit using fpr under the conditions tested, so we moved forward using the optimized DSD conditions (with cofactor regeneration for formate as a substrate) without fpr for further testing of ReForm. With formate as a substrate, final reaction conditions were 20 mM sodium phosphate buffer pH 7.4, 2 mM ATP, 4 mM NADH, 10 mM MgCl_2_, 1 mM CoA, 50 µM TPP, 2 mM glyoxylate, 100 mM ^13^C sodium formate, 10 mM polyphosphate, 5 U/mL pyrophosphatase (Sigma-Aldrich I5907), 2 µM acs, 5 µM acr_f_, 5 µM acr_g_, 20 µM oxc, 20 µM pk, 0.01 µM pta, 0.5 µM mls, 2 µM ppk1, 2 µM ppk2, and 1 µM fdh. The same conditions were used with formate derived electrochemically from CO_2_ as a substrate, with the exception of a final concentration of 8 mM formate (^12^C) being used. Given formate was produced electrochemically at a concentration of 42 mM, this comprised ∼20% v/v of the reaction.

With formaldehyde as a substrate, final reaction conditions were 20 mM sodium phosphate buffer pH 7.4, 2.19 mM NAD+, 10 mM MgCl_2_, 0.78 mM CoA, 50 µM TPP, 2 mM glyoxylate, 20 mM formaldehyde, 5 µM acr_f_, 5 µM acr_g_, 15.5 µM oxc, 20 µM pk, 0.384 µM pta, and 0.5 µM mls. Reactions were performed in triplicate at 20 µL with time points as follows (hours): 0, 0.17, 0.33, 0.5, 0.75, 1, 1.5, 2, 3, 6, 10, 18, 24, 36, 48. The same reaction conditions were used with methanol as a substrate with the following exceptions. Final reactions additionally contained 300 U/mL catalase (Sigma-Aldrich C1345), 1 U/mL alcohol oxidase (Sigma-Aldrich A2404), and 20 mM methanol.

### Calculation of pathway metrics

All rates (steady-state productivity, specific productivity, and CO_2_ fixation rate) were calculated using the optimized ReForm pathway results shown in **Fig. 4C**, where formate is derived from the electrochemical reduction of CO_2_. Malate (and equivalently acetyl-CoA) synthesis rates can be aggressively calculated from the steady-state portion of the pathway between 0 and 6 hours (280 µM malate in 6 hours yields) or more conservatively by taking the highest yield at 24 hours (607 µM malate in 24 hours). We additionally consider the spatially and temporally separate electrochemical module that reduces CO_2_ to formate, which ran for 1 hour total (280 µM malate in 7 hours yields 40.1 uM hr^-1^ or 668 nM min^-1^ and 607 µM malate in 25 hours yields 24.3 uM hr^-1^ or 404 nM min^-1^). We report 40.1 uM hr^-1^ as the steady-state productivity in **Table S1**. The total enzyme loading for this reaction was 4.073 mg protein (**Table S9**), which gives a specific productivity of 9.8 uM hr^-1^ mg^-1^ protein. The specific productivity of CO_2_ equivalents assimilated was calculated using this value and considering that for every molecule of malate (and acetyl-CoA) produced, 2 molecules of CO_2_ equivalents (formate) are assimilated. We also include these values calculated for other state-of-the-art synthetic CO_2_ fixing pathways (the CETCH cycle and the THETA cycle). It can be difficult to quantitatively compare these values, but we feel it is important to wholistically compare how each pathway functions and their inherent advantages and disadvantages.

### Catalyst preparation and electrochemical reduction of CO_2_

All chemicals used for electrolytes, catalyst synthesis, and electrode preparation, including iridium(III) chloride hydrate (IrCl_3_·xH_2_O, 99.9%), Tin (IV) oxide (SnO_2_, nanopowder, ≤100 nm) and Potassium bicarbonate (KHCO_3_, 99.7%) were purchased from Sigma Aldrich and used without any further treatments. Nafion 117 membrane and titanium felt were purchased from the Fuel Cell Store. In a typical procedure, 20 mg SnO_2_, and 60 μL Nafion perfluorinated resin solution were stirred and sonicated in 4 mL absolute ethanol to form the catalyst ink. Then, 4 mL of the ink was sprayed onto the gas diffusion layer (Freudenberg H23C3) with a SnO_2_ loading of 1 mg cm^-2^. All electrochemical experiments were performed in an MEA electrolyzer (SKU: 68732; Dioxide Materials), with a gasket to control the active area of 1 cm^2^ accessed with a serpentine channel, unless otherwise specified. A proton exchange membrane (Nafion 117) was sandwiched between the cathode and Titanium felt-supported iridium oxide (IrO_2_/Ti) anode. The IrO_2_/Ti anode was fabricated according to a reported method^54^ with slight modifications. Unless otherwise specified, the anode was circulated with 0.5 M KHCO_3_ electrolyte at a rate of 10 mL/min with a peristaltic pump and with silicone Shore A50 tubing. 40 sccm of humidified CO_2_ was fed into the cathode using an accurate mass flow controller. The cathodic product is collected in 10 mL of DI water. The volume of anolyte used was 20 ml. Electrochemical measurements were carried out using an Autolab PGSTAT204 in an amperostatic mode and a current booster (10 A). Unless otherwise stated, at the end of 1 h of electrolysis, a sample of cathodic product and a sample of anolyte were extracted for liquid product analysis. Cathodic product and anolyte sample were identified by ^1^H NMR spectroscopy (600 MHz Agilent DD2 NMR Spectrometer) with DMSO internal standard using water suppression techniques.

### Data collection and analysis

All statistical information provided in this manuscript is derived from *n*L=L3 independent experiments unless otherwise noted in the text or figure legends. Error bars represent 1 s.d. of the mean derived from these experiments. Data analysis and figure generation were conducted using Excel Version 2304, ChimeraX Version 1.5, GraphPad Prism Version 9.5.0, and Python 3.9. Data collected on the BioTek Synergy H1 Microplate Reader were analyzed using Gen5 Version 2.09.2.

## Acknowledgements

Molecular graphics and analyses performed with UCSF ChimeraX, developed by the Resource for Biocomputing, Visualization, and Informatics at the University of California, San Francisco, with support from National Institutes of Health R01-GM129325 and the Office of Cyber Infrastructure and Computational Biology, National Institute of Allergy and Infectious Diseases. This work made use of the IMSERC MS facility at Northwestern University, which has received support from the Soft and Hybrid Nanotechnology Experimental (SHyNE) Resource (NSF ECCS-2025633), the State of Illinois, and the International Institute for Nanotechnology (IIN). We thank Stanford University Cryo-EM center (cEMc) and particularly Bharti Singal for providing support for Cryo-EM grid preparation, data collection, processing and structure determination pipeline. We thank Fernando “Ralph” Tobias for his help in developing analytical methods for malate detection and Kosuke Seki for his help in gathering intact protein mass spec data on the deacetylated acyl-CoA synthetases. We also thank Jonathan W. Bogart for conversations regarding this work.

## Funding

Department of Energy (DE-SC0023278)

NSF GRFP

## Author contributions

Conceptualization: ASK, BV, GML, MCJ

Methodology: BV, GML

Investigation: AG, CT, GML, RL

Cryo-EM: BS

Supervision: ASK, MCJ, EHS

Funding acquisition: ASK, MCJ, EHS

Writing: ASK, GML, MCJ

## Data and materials availability

All data are available in the main text or the supplementary materials. Atomic structures reported in this paper are deposited to the Protein Data Bank under accession codes 9CD3 and 9CD4. The cryo-EM data were deposited to the Electron Microscopy Data Bank under EMD-45461 and EMD-45462.

